# Bcl-xL dynamics and cancer-associated mutations under the lens of protein structure network and biomolecular simulations

**DOI:** 10.1101/574699

**Authors:** Valentina Sora, Elena Papaleo

**Affiliations:** Computational Biology Laboratory, Danish Cancer Society Research Center, Strandboulevarden 49, 2100, Copenhagen, Denmark; Translational Disease Systems Biology, Faculty of Health and Medical Sciences, Novo Nordisk Foundation Center for Protein Research University of Copenhagen, Copenhagen, Denmark

**Keywords:** PUMA, molecular dynamics, p53, apoptosis, cancer mutations, allostery, long-range communication

## Abstract

Understanding the finely orchestrated interactions leading to or preventing programmed cell death (apoptosis) is of utmost importance in cancer research since the failure of these systems could eventually lead to the onset of the disease. In this regard, the maintenance of a delicate balance between promoters and inhibitors of mitochondrial apoptosis is crucial, as demonstrated by the interplay among the Bcl-2 family members. Particularly, Bcl-x_L_ is a target of interest due to its forefront role of its dysfunctions in cancer development. Bcl-x_L_ prevents apoptosis by binding both the pro-apoptotic BH3-only proteins, as PUMA, and noncanonical partners such as p53 at different sites. An allosteric communication between the BH3-only proteins binding pocket and the p53 binding site has been postulated and supported by NMR and other biophysical data, mediating the release of p53 from Bcl-x_L_ upon PUMA binding. The molecular details, especially at the residue level, of this mechanism remain unclear. In this work, we investigated the distal communication between these two sites in both Bcl-x_L_ in its free state and bound to PUMA, and we evaluated how missense mutations of Bcl-x_L_ found in cancer samples might impair the communication and thus the allosteric mechanism. We employed all-atom explicit solvent microsecond molecular dynamics simulations analyzed through a Protein Structure Network approach and integrated with calculations of changes in free energies upon cancer-related mutations identified by genomics studies. We found a subset of candidate residues responsible for both maintaining protein stability and for conveying structural information between the two binding sites and hypothesized possible communication routes between specific residues at both sites.

## Introduction

B-cell lymphoma extra-large (Bcl-x_L_) is one of the best-known proteins of the Bcl-2 family, whose members have been extensively studied over the past decades due to their role in apoptosis regulation, tumorigenesis and cellular responses to anti-cancer therapies ^1^.

The members of this family can be subdivided in three classes, according both to their role in apoptosis and to the different functional domains shared with the parent Bcl-2. The first class includes proteins containing all the four *B*cl2-*H*omology domains (named BH1-2-3-4) and eliciting an anti-apoptotic response, whilst the second consists of pro-apoptotic proteins containing only the BH3 domain (BH3-only proteins) and the third is formed by proteins including all the BH domains but performing a pro-apoptotic function ^2–4^.

Bcl-x_L_ belongs to the first class, and structurally consists of eight α-helices connected by loops of different lengths (**Fig. 1**) and a transmembrane C-terminal region recently solved ^5^. Bcl-x_L_ carries out its anti-apoptotic function by binding a variety of different partners, such as p53 ^6,7^ or other Bcl-2-family members, including many BH3-only proteins such as PUMA ^3,8^.

**Fig.1.**
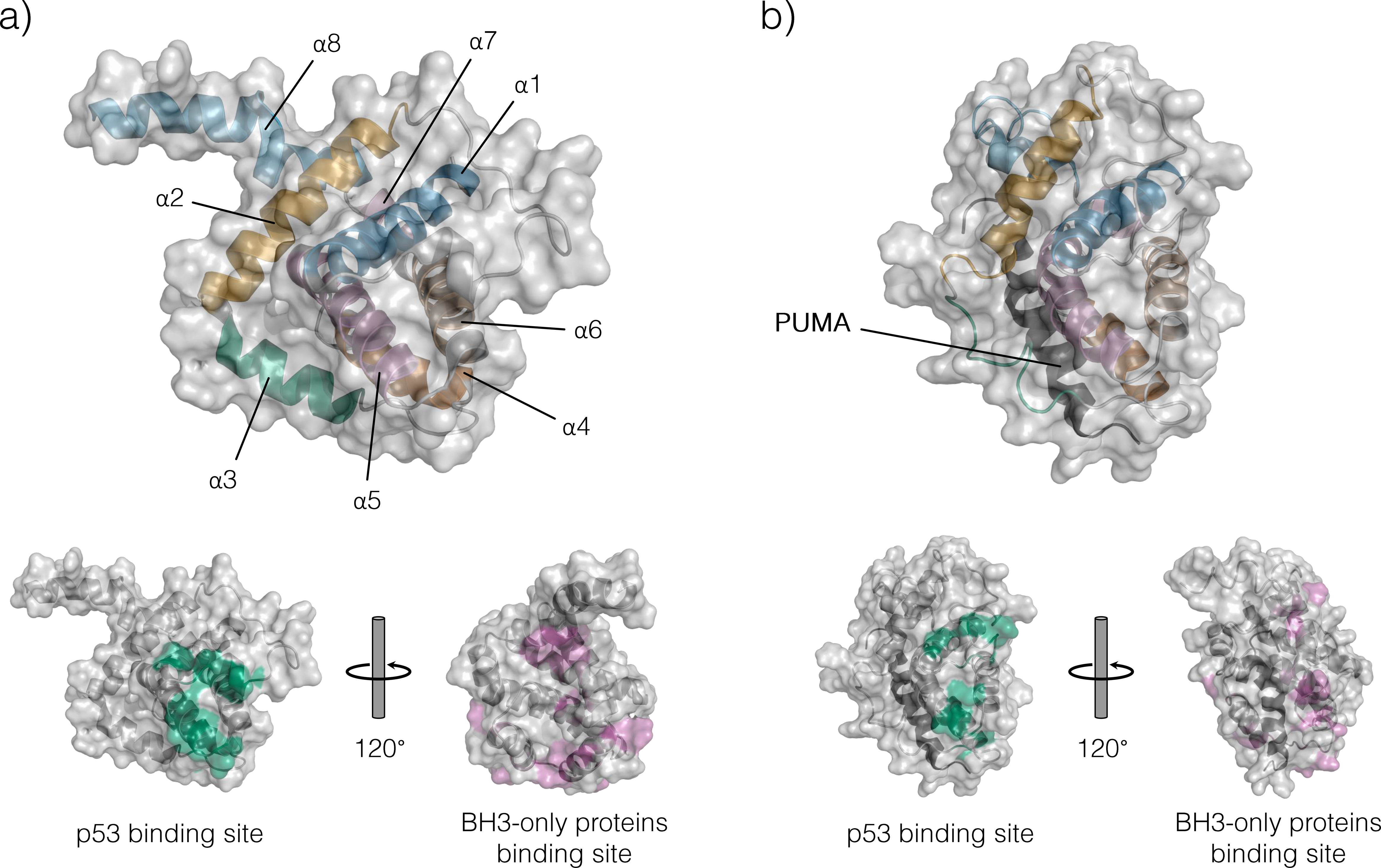
Structure of Bcl-x_L_ free and of the Bcl-x_L_-PUMA complex a) Structure of Bcl-x_L_, missing the loop α1-α2, from the PDB structure 2LPC. The eight helices are colored and labelled. b) Structure of Bcl-x_L_ bound to PUMA from the PDB structure 2M04. The eight helices are color-coded as in the free structure, and PUMA is shown in black.

It has been demonstrated that the interaction of Bcl-x_L_, with the BH3-only proteins could drive long-range structural perturbations within Bcl-x_L_, such as the one observed at the p53 binding site upon the binding of PUMA into the Bcl-x_L_ binding pocket for BH3 domains ^9,10^. On the other hand, this allosteric communication may work both ways, determining conformational changes into the BH3-only binding pocket when p53 is bound ^11^.

Allosteric mechanisms involve the transmission of local structural perturbations in a cascade of couplings of motions or collisional contacts that can be described as paths of communication from one site to a distal one within the protein structure ^12–16^. These mechanisms can involve large conformational rearrangements or subtle changes in dynamics and local conformations ^17,18^, and allostery is likely to be an intrinsic feature of any protein ^17,19^. In this context, it can be described as a shift in the distribution of pre-existing states that are present in both the free and bound conformational ensemble of the protein ^20–22^.

The network of residue contacts and the intrinsic dynamics of a protein are thus key components to understand long-range communication and the related molecular and functional mechanisms. One useful framework is the so-called Protein Structure Network (PSN) paradigm, which has been extensively used to describe the structure, topology, and dynamics of proteins ^23–28^. In the PSN, intramolecular noncovalent interactions between residues pairs in a protein can be collectively represented as a network ^28–30^.

Nevertheless, protein structures are not static entities. Indeed, they undergo a plethora of different motions and conformational changes and are better described as an ensemble of different conformational states in a dynamic equilibrium ^22,31–36^ that can be perturbed by post-translational modifications, mutations or interactions with binding partners ^33,37–39^. These biologically meaningful dynamic processes can be investigated by computational methods such as molecular dynamics (MD) simulations, which allow an atom level description of the protein conformations ^40–42^. The integration of PSN approaches and MD itself proved useful in the last decade to shed light over mechanisms of allosteric communication paths between residues in proteins ^13,26,43–55^. The study of protein ensembles from the PSN perspective also revealed to be useful in identifying paths of long-range communication that are only activated upon perturbations within the ensemble (i.e. ligand binding or mutations), despite being present also in the unperturbed system. ^28,56,57^

In light of the above observation, we applied a MD-PSN pipeline to the study of the conformational ensemble of Bcl-x_L_. At first, we aimed to verify if the MD-PSN approach can describe the allosteric mechanisms induced by PUMA binding. Then, we investigated how the distal structural communication is propagated and to pinpoint the key residues responsible for the communication between the p53 and the BH3 binding sites of Bcl-x_L_, so that a more profound understanding of this fundamental allosteric mechanism can be achieved. Furthermore, we evaluated if the paths of structural communication between these two distal binding regions were an intrinsic feature of Bcl-x_L_ and, as a such, if they were present in Bcl-x_L_ even in the free state and then reinforced or (de)activated upon binding of the BH3-only PUMA protein.

## RESULTS

### The interaction with PUMA causes the loss of edges in the p53 binding sites

We collected microsecond all-atom MD simulations for Bcl-x_L_ in its free and bound state to the BH3-only protein PUMA and used this ensemble to infer a contact-based protein structure network of the two forms of the protein.

We extracted the edges common to all the PSNs (i.e., the three Bcl-x_L_ free simulations and the simulation of the complex Bcl-x_L_-PUMA) and compared them to the ones only common to the three PSNs of the Bcl-x_L_ free MD ensembles to appreciate the effect induced by the binding of PUMA on Bcl-x_L_ structure and dynamics (Figure 2).

**Fig.2.**
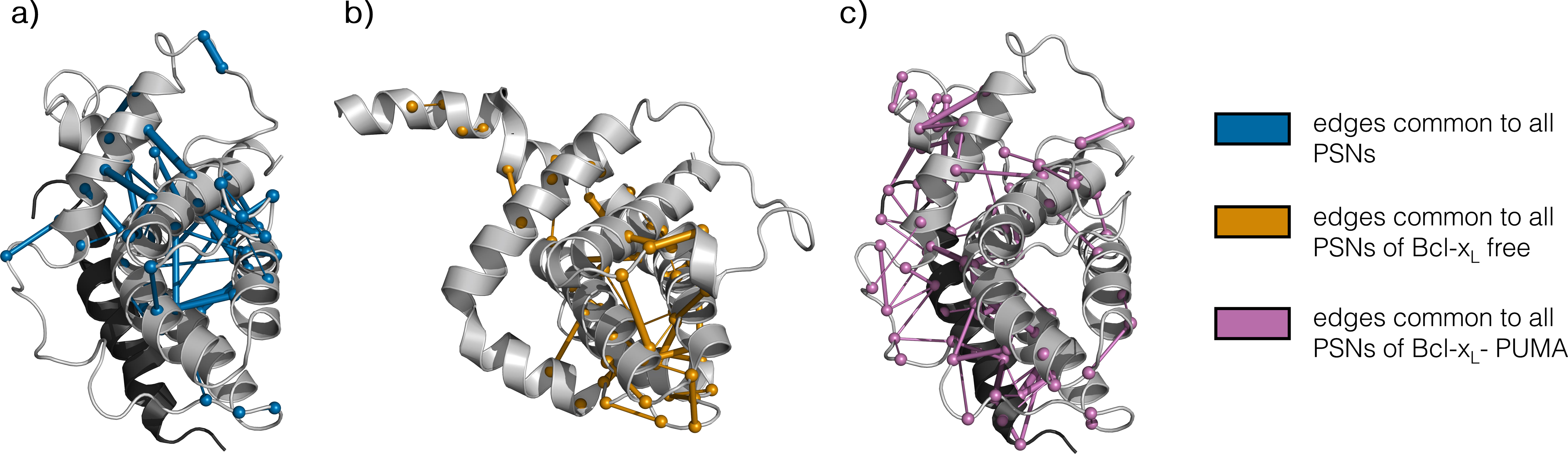
Common edges Common edges for all PSNs (a) for all Bcl-x_L_ free PSNs (b) and for Bcl-x_L_-PUMA (c). Edges are displayed as sticks and mapped on the PDB structure 2M04 for the Bcl-x_L_-PUMA complex and on 2LPC for Bcl-x_L_ free, respectively. The thickness of each edge is proportional to its persistence. The Cα of residues connected by edges are displayed as balls.

We noticed that the majority of the edges conserved in all PSNs were located in the core of the protein with a high persistence, suggesting a major architectural role for these interactions. We also noted that all the interactions with high occurrence in the Bcl-x_L_ free MD ensembles between the loops ◻1-◻2, ◻3-◻4 and the N-terminal portion of ◻4 disappeared when PUMA was bound to the protein (Figure 2). This region largely overlap with the p53 binding site of Bcl-x_L_ (Figure 1B and see below for more details), which is located approximately 21 Å apart from the BH3 binding site. The massive loss of interactions at the p53 binding site upon PUMA interaction into the BH3-binding pocket of Bcl-x_L_ may indicate a structural perturbation transmitted long range from one site to the other.

### Intrinsic and ligand-induced hub as predictive of interfaces for recruitment of binding partners of Bcl-x_L_

Hubs are the residues with highest connectivity in a network, i.e. nodes connected by more than three edges in a PSN ^26^. They can play a role in both protein structural stability and protein function or allow a proper flux of information between distal sites ^13,25,28^. Particularly, proteins are known to be built of a significant number of strongly and weakly interacting hub residues stabilizing the tertiary structure by providing resilience against random mutations ^20,58–60^.

We identified 14 hubs conserved in all the MD replicates of Bcl-x_L_ in its free state, which are mostly located in the core ◻-helices. Five of them were also present in the MD ensemble of Bcl-x_L_ in complex with PUMA (Fig. 3A), whilst the others were specific of the unbound Bcl-x_L_ (Fig. 3B). Of particular interest are F146, experimentally observed as important for the binding of the BH3-only proteins to Bcl-xL ^11,61^ and C151, whose mutation C151Y features a high pathogenicity score according to the REVEL ensemble predictor ^62^ (Table S1).

**Fig. 3.**
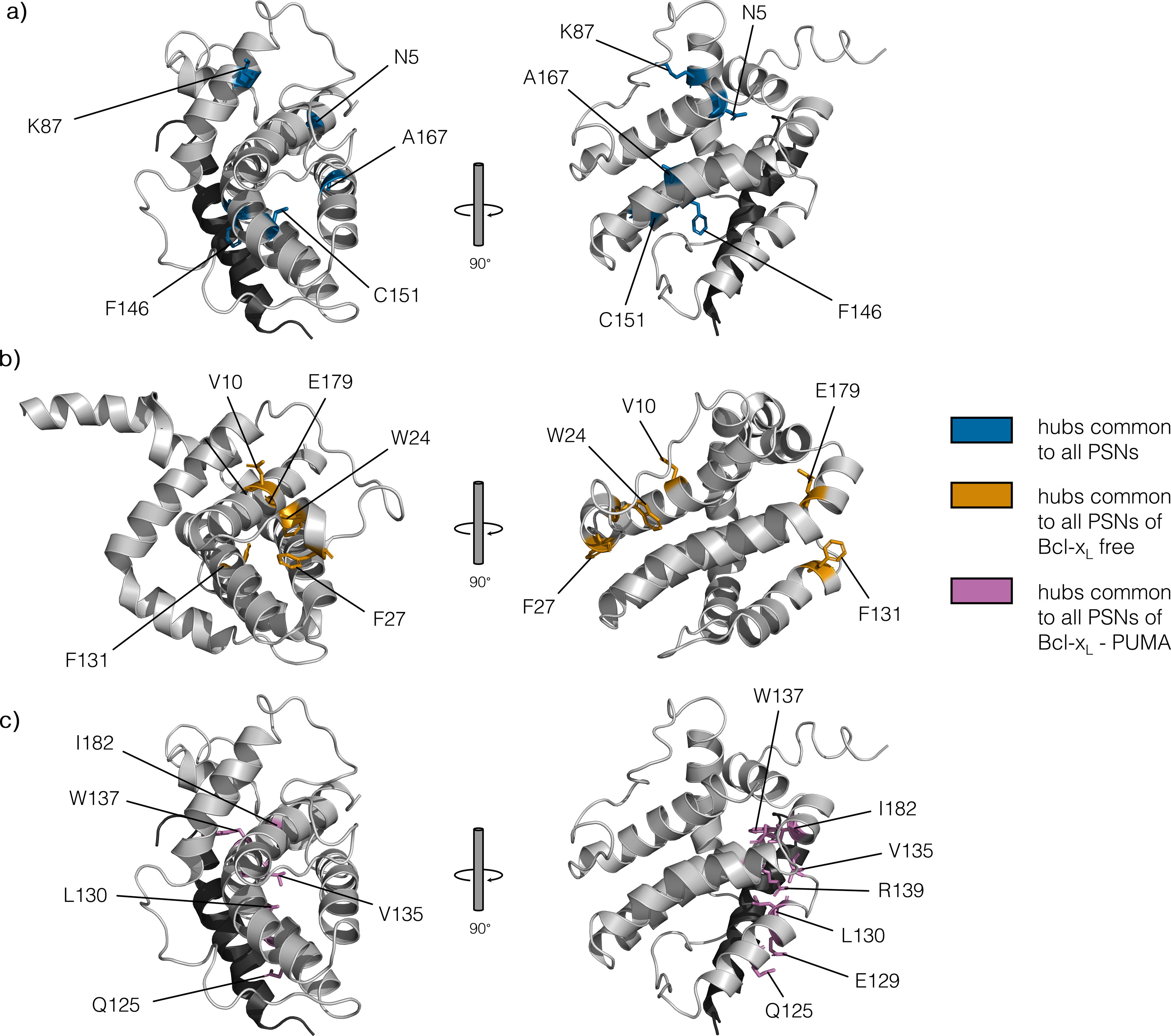
Common hubs Common hubs for all PSNs (a) for all Bcl-x_L_ free PSNs (b) and for Bcl-x_L_-PUMA (c). Hub nodes are shown as sticks. Bcl-x_L_ is showed in grey, PUMA in black.

The interaction with PUMA modified the Bcl-x_L_ PSN toward an enrichment of hubs located into the BH3-only proteins binding pocket which were absent in the free state of the protein (i.e., A104, A119, S122, Q125, E129, L130, N136, R139, A142, L194). Hagn and coworkers ^11^ have reported some of these residues undergoing significant structural perturbations when Bcl-x_L_ binds p53 at a distal site with respect to the BH3-binding groove using NMR chemical shift perturbation and Paramagnetic Relaxation Enhancement (PRE) measurements. In particular, they reported as key residues sensitive to the allosteric effect induced by p53-binding A104, A119, S122, Q125, A142, L194 ^11^, enforcing the notion that the binding of p53 influences the binding mechanism of the BH3-only PUMA protein. Moreover, the work of Campbell and coworkers ^61^, using a cell free split-luciferase assay, identified four of the PUMA-induced hubs (E129, L130, N136, R139) as crucial residues for the binding of several BH3-only proteins, including PUMA. Two of these residues, L130 and R139, are identified as hubs only in the ensembles of the Bcl-x_L_-PUMA complex, reinforcing the hypothesis of a prominent role in mediating the binding process.

To investigate how the mutations in the five conserved hubs and in those specific of Bcl-x_L_ in free state and of the Bcl-x_L_-PUMA complex might impact the protein stability or binding free energy, we calculated the variation of the ΔΔG upon all possible missense variations at each site for both the free Bcl-x_L_ NMR ensemble (Supplementary File S1) or the NMR ensemble of the complex with PUMA (Supplementary File S2). As shown in Fig. 4A, mutations in C151, S154 and V163 were predicted to significantly destabilize the protein structure but with minor impact of binding.

**Fig 4.**
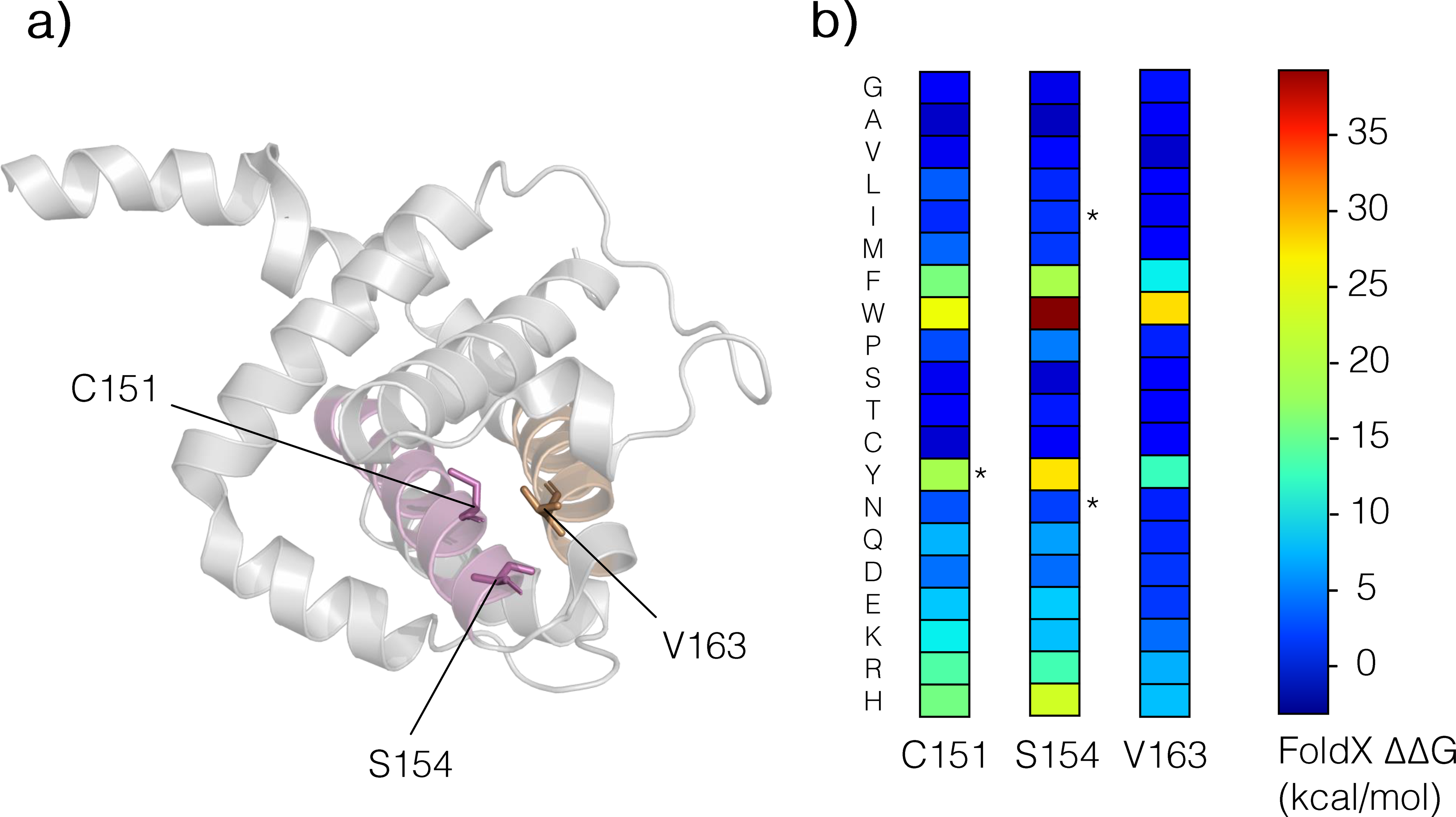
Mutational hotspots in Bcl-x_L_ Hotspot residues are displayed as sticks and labelled, color-coded according to the helix they belong (α5 is pink, α6 is orange). b) Structural impact of mutations at hotspots. The results from the FoldX scan are shown as a heatmap, with each mutation color-coded according to the value of the calculated ΔΔG with respect to the wild-type variant.

Our PSN results, together with the experimental data from NMR and biochemical assays mentioned above, suggest that the PUMA-induced hub residues are predictive of the key residues for the interaction with the BH3-only proteins.

### Identification of paths of long-range communication between the p53 binding site and the BH3-only proteins binding pocket

We selected the pairs of residues for shortest paths calculation on the basis of the work of Hagn and coworkers ^11^, who experimentally identified the residues constituting the binding interface between p53 and Bcl-x_L_ and postulated the existence of a distal communication between the p53-binding site and the BH3-only proteins binding groove of Bcl-x_L_ (Fig. 5A). The interaction of p53 with Bcl-x_L_ leads to a conformational change in the pocket, conversely resulting in the displacement of the BH3-protein possibly bound ^11^ and to the initiation of the apoptotic process. This work has been corroborated by Follis and coworkers ^9^, who described a long-range communication between the two binding sites resulting from the binding of PUMA, a pro-apoptotic BH3-only protein, to the BH3-proteins binding pocket. Indeed, they noted that the formation of the Bcl-x_L_-PUMA complex was accompanied by the displacement of p53 from Bcl-x_L_ and postulated a two-way long-range structural route driving this phenomenon ^9^.

**Fig.5.**
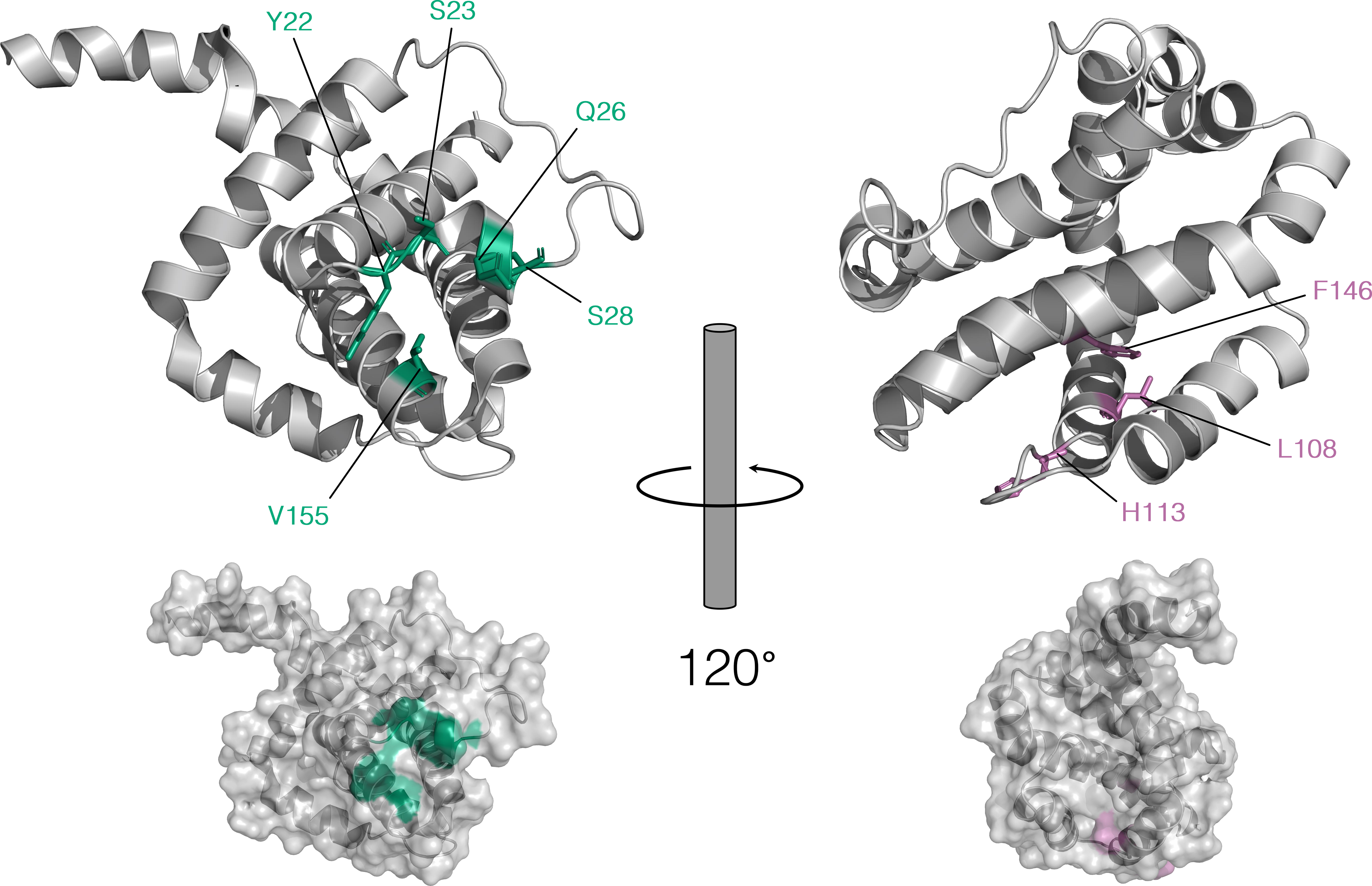
BH3-only proteins and p53 binding sites The p53 binding site is colored in green, whilst the BH3-only proteins binding pocket is shown in pink. The residues crucial for the binding process and for which at least a shortest path exists are shown as sticks and labelled.

Therefore, we calculated the shortest paths between the residues determined as fundamental for the binding of p53 (S18, Y22, S23, Q26, S28, I114, V155, D156, E158) and those in the BH3-only proteins binding pocket (A93, F97, R102, A104, S106, L108, Q111, A119, S122, Q125, F131, A142, F146, L194, Y195) showing perturbed chemical shifts upon p53 binding according to Hagn’s work (Fig.5B). We also included an additional residue identified by Kriwacri’s group ^9^ as crucial for PUMA binding, H113. We aimed: i) to verify if any communication path exists that may transmit the perturbation to the p53 binding site, ii) if these paths are pre-existing routes in the free ensemble of Bcl-x_L_ in solution and iii) what are the key residues involved in the allosteric communication.

We observed that shortest paths exist in all simulation replicates of Bcl-x_L_ free for the following residue pairs (the first element of each pair is a residue belonging to the p53 binding site, the second lies into the BH3-only proteins binding pocket): Y22-L108, Y22-H113, Y22-F146, S23-L108, S23-H113, S23-F146, Q26-L108, Q26-H113, Q26-F146, S28-L108, S28-H113, S28-F146, V155-L108, V155-H113, V155-F146.

We noted that almost all paths having S28 as first residue shared three residues of the path (V10, A167, L13) but none of these residues appear in other paths, suggesting a specificity of those residues in conveying structural information from S28 to the BH3-only proteins binding pocket. The only exception is path S28-H113 of replicate 3, in which another route is identified, lacking both V167 and L13. V10, the only one ubiquitously present, has been reported in COSMIC as cancer-related ^63^.

Similarly, all paths reaching F146 share the three terminal residues (C151, I166, L150). We observed that the same residues are also part of the terminal portion of all paths ending in L108, meaning that L108 and F146 share part of their communication route with the p53 binding site. Moreover, excluding from the paths ending either in L108 or F146 the two ones having S28 as extreme, we noticed that the number of intermediate shared residues increased (P27, V123, C151, I166, L150), thus making these paths almost perfectly superposable with one another. Our results strongly indicates that there are pre-existing communication routes between the p53- and the BH3-binding sites of Bcl-x_L_. We also identified a group of residues that are crucial players in the modulation of the long-range communication between the two binding sites, which could be sensitive sites for mutations to disrupt the allosteric mechanism.

We calculated the same paths for the ensemble of the Bcl-x_L_-PUMA complex, and we observed that a path existed only for the pairs Y22-L108, Y22-F146, S28-L108, S28-F146, V155-L108, V155-F146. We found that paths ending in F146 were all equally long and shared 5 out of 6 intermediate residues, thus portraying a shared communication route from residues belonging to the p53 binding site to F146. Considering their counterparts in the replicates of Bcl-x_L_ free, paths in the Bcl-x_L_-PUMA complex are shorter and share only three residues (C151, I166, L150) with the corresponding ones in the Bcl-x_L_ free ensembles. The only exceptions are paths ending in L108 in the Bcl-x_L_-PUMA ensemble, they are considerably longer with respect to their analogous in the Bcl-x_L_ free ensembles, encompassing 20 residues instead of 7-10 Furthermore, the topological connections between these common residues are different. Therefore, despite displaying few similarities, the routes connecting the p53 binding site to F146 when Bcl-x_L_ is in its free state and when PUMA is bound are distinct, zig-zagging among different helices in the core of the protein. Thus, we can conclude that the binding of PUMA induces a weakening of the pre-existing routes.

### C151, S154 and V163 are involved both in protein stability and in long-range communication between the p53 binding site and the BH3-only proteins binding pocket

C151 and S154 are both localized on α5, with their side chains protruding toward α6. On the other hand, V163 is localized on α6 and their side chains point toward α5 (Fig. 4, A).

C151, identified as hub in all the ensembles, is ubiquitous in paths ending in either L108 or F146 and is also present in the path S28-H113, thus appearing in eleven out of fifteen identified paths in the Bcl-x_L_ free ensembles, and in all six paths observed in the Bcl-x_L_-PUMA ensemble. This finding indicates a possible front role in conveying structural information between the BH3-only proteins binding pocket and the p53 binding site. The structural relevance of C151 is further supported by the results of the in-silico mutational scan displaying a wide range of mutations of C151 predicted to impact the stability of Bcl-xL (Fig. 4, B). Particularly, C151Y is predicted as strongly destabilizing (ΔΔG around 20 kcal/mol), and has also been predicted as highly pathogenic by its REVEL score (0.567) (Table S1).

S154 is also a hub in all the networks of Bcl-x_L_ in free state, and is found in all paths having H113 as extreme. As for C151, our mutational scan predicted nearly half of its possible mutations as highly destabilizing. Interestingly, mutations S154I and S154N, found in cancer samples but predicted as non-pathogenic by their REVEL scores, are predicted as weakly impacting on the protein stability by FoldX. Therefore, our scan corroborates the hypothesis that these particular mutations have only a weak impact on Bcl-x_L_ and its functionality, but at the same time suggests that other mutations of S154 may influence significantly the protein stability.

Both C151 and S154 have been also proposed as core structural residues by a comparative study on the conservation of sequence and structure among the Bcl-2 family members ^64^.

V163 is alsoidentified as hub and it is present in 13 out of 15 identified paths in the Bcl-x_L_ free ensembles. As for C151 and S154, its mutations are predicted as highly destabilizing the protein fold.

In summary, the identification of C151, S154 and V163 as strong hubs, the significant impact of their mutations on protein stability as predicted by the mutational scan and their localization into the protein core suggest a major involvement in both maintaining the protein core architecture and in transferring structural perturbations from the p53 binding site to the BH3-only proteins binding pocket and vice versa.

### The communication route between the p53 binding site and the BH3-only binding pocket is interrupted when PUMA is bound

In contrast to what we observed for the ensembles of Bcl-x_L_ in free state, we found no paths having H113 as extreme in one of the replicates of the Bcl-x_L_-PUMA complex, whilst in the other replicate no paths at all were found between the selected residues. Furthermore, H113 forms a connected component on its own in the former network, meaning that it is isolated from the rest of the protein, whilst in latter one all residues belonging to the BH3-only proteins binding site cluster in one connected component, still isolated from the rest of the protein. On the contrary, H113 belongs to the most populated connected component in all Bcl-x_L_ free ensembles thanks to the edge formed with T115. Being the only edge connecting H113 to the rest of the component, it is also part of all paths of communication with the p53 binding site. It is present in nearly half of the structures in all the Bcl-x_L_ free ensembles (~46%), meaning that the communication route between H113 and the p53 binding site is not always active. This suggests that a binding phenomenon perturbing H113 may shift the equilibrium towards a permanent activation or deactivation, as we observed when PUMA was bound. Indeed, in this case the edge was present in 3.1% of the structures, being thus removed in the final PSN (built with cut-off 20%, as explained in the Methods section). This implies an increased distance between H113 and T115 which interrupts the communication, as also shown by the calculation of the distance over time in all the simulations (Fig. 6).

**Fig. 6.**
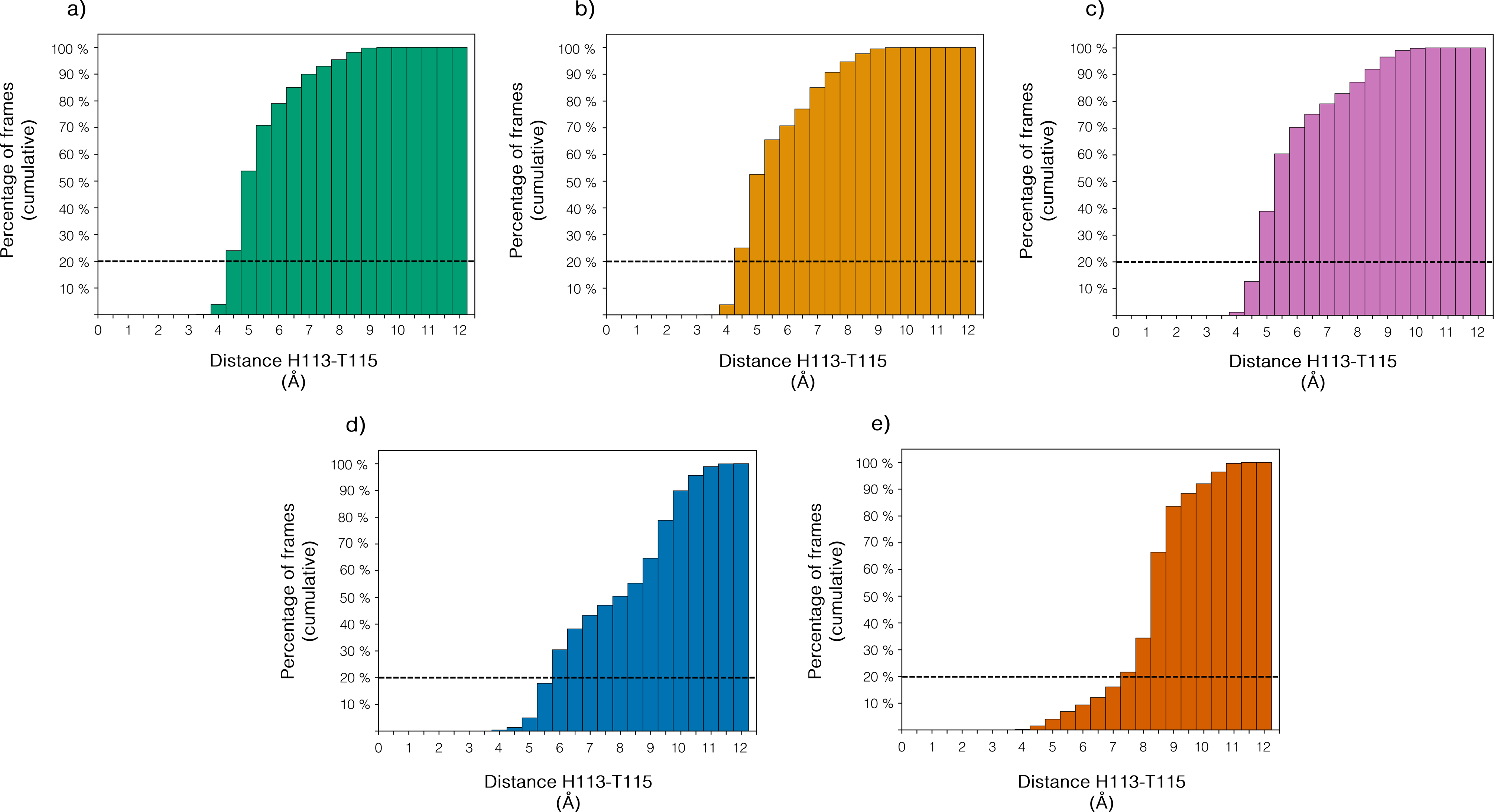
Distribution of the H113-T115 distance within the PSNs Cumulative histograms showing the distribution of the H113-T115 distance within the five PSNs. The three replicas of Bcl-x_L_ free are displayed in green, orange, pink, whilst the PSNs for the Bcl-x_L_-PUMA complex are shown in blue and red. The horizontal dashed lines indicate the persistence cut-off in the PSNs, meaning that if the distance is lower than 5.125 Å (distance cut-off in the PSNs) in less than 20% of frames, no edge is present between the two residues.

Since PUMA has been the only BH3-only protein identified so far as capable of triggering the release of p53 from Bcl-x_L_ and H113 has been postulated as crucial for the interaction of PUMA with Bcl-x_L_ itself [23340338], the sudden disruption of the paths between H113 and the p53 binding site upon PUMA binding may indicate the loss of an internal structural communication that elicits the release of p53 (Fig. 7).

**Fig. 7.**
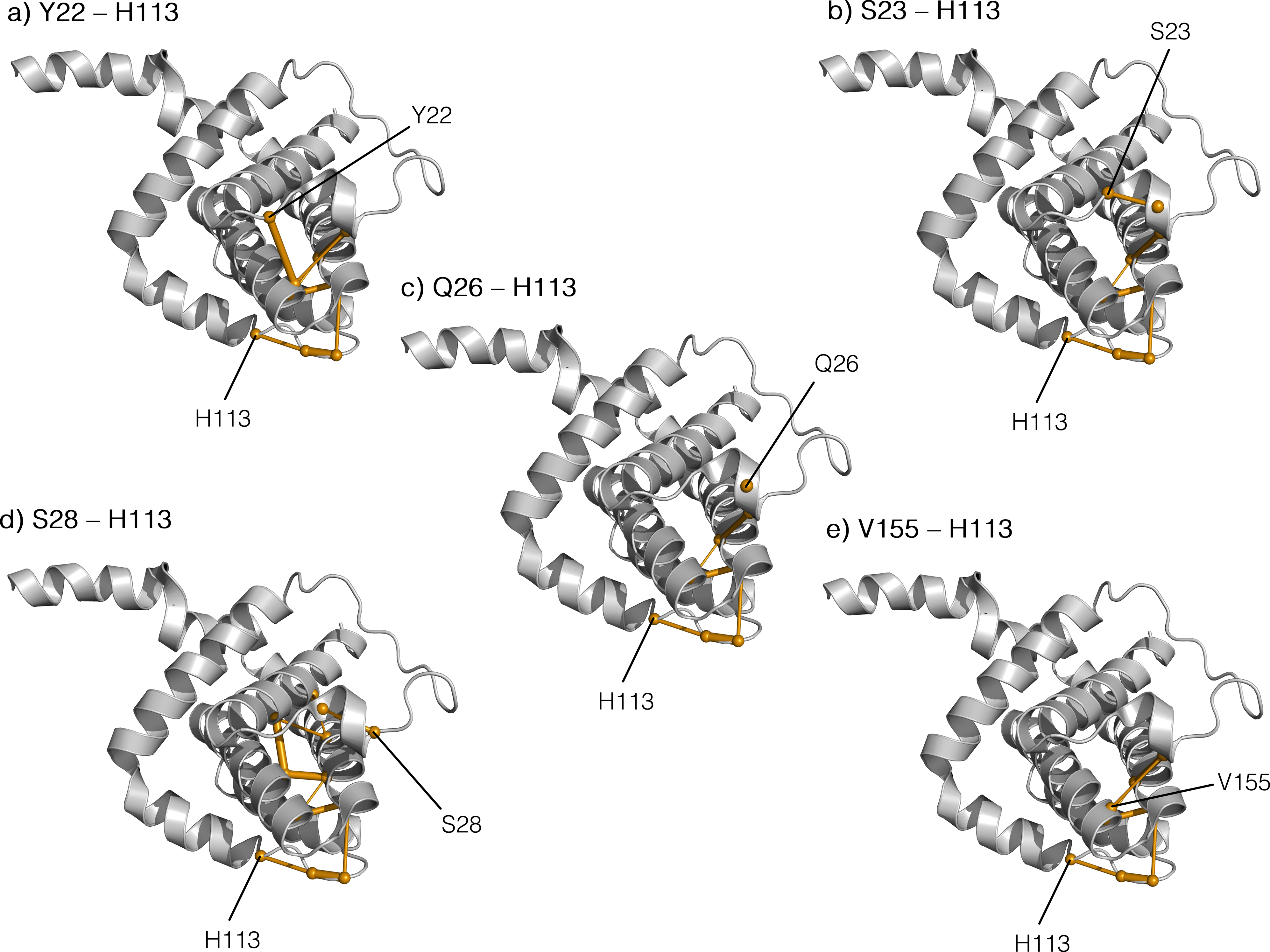
Paths from the p53 binding site to H113 in Bcl-x_L_ free Paths connecting the p53 binding site to H113. They are present in all the PSNs of Bcl-x_L_ free, but for clarity purposes only those in the PSN computed from the TIP3P+capping ensemble are displayed here. The thickness of each edge is proportional to its persistence, and the Cα of residues belonging to each path are displayed as balls.

## METHODS

### Molecular Dynamics simulations

We carried out three replicates of one-microsecond all-atom explicit solvent molecular dynamics (MD) simulations with *GROMACS* version 5^65^ with the CHARMM22*^66^ force field for Bcl-x_L_ free and the Bcl-x_L_/PUMA complex.

For Bcl-x_L_ free, we employed the first model of the NMR-derived ensemble of Bcl-x_L_ in the PDB entry 2LPC^67^ as starting structure for the MD simulations, whilst for the Bcl-x_L_/PUMA complex we used the first model of the PDB entry 2M04 ^9^. The Bcl-x_L_ free structure spans residues 1-44 and 85-209 of Bcl-x_L_ (when compared to the UniProt sequence, isoform 1), missing a highly flexible loop between the first two helices and the membrane-inserted helix at the C-terminus. Only one structure of full-length Bcl-x_L_ was available in the PDB at the time of writing (NMR ensemble, PDB entry 1LXL^68^), but its predicted resolution according to ResProx was too high (4.932 Å) to be considered as starting structure for our simulations, and the flexible loop was too long to be reliably modeled. Furthermore, many experimental studies have been carried out on the deleted variant of Bcl-x_L_, which is also fully functional in cellular experiments ^68,69^. Hence, in order to maintain a good balance between sequence coverage and resolution, we selected the first model of the NMR ensemble 2LPC as starting structure. We first removed the HIS tag at the C-terminus of 2LPC, therefore retaining residues 1:169 of the PDB structure.

We collected three replicates for the free form of Bcl-x_L_ and two replicates of the Bcl-x_L_-PUMA complex to ensure the reproducibility of our results, testing the impact of changing either the solvent model (from TIP3P to TIPS3P) or the N- and C-terminal (from capped to charged).

In one of the replicates of Bcl-x_L_ in free state and in one of those of the complex, the starting structure was capped at both termini to avoid effects due to terminal partial charges. The N-terminus and C-terminus were made neutral by removing a proton and by adding an amide capping group, respectively. In particular, the choice of amidating the C-terminus was justified by the actual continuation of the backbone beyond our C-terminus in the full-length protein.

The starting structure was soaked in a dodecahedral box of water molecules at 150 mM [NaCl], using the TIP3P solvent model ^70^. The edges of the box were all at least 20 Å distant from the protein atoms in all the Bcl-x_L_ free simulations, and 35 Å and 25 Å in the two replicates of the Bcl-x_L_/PUMA complex, respectively. We simulated each system under periodic boundary conditions.

We equilibrated the systems according to a protocol previously applied to other cases of study ^50,51^, and we performed productive MD simulations in the canonical ensemble at 300 K using velocity rescaling with a stochastic term. We used the LINCS algorithm to constrain the heavy atom bonds ^71^ to allow the usage of a 2 fs timestep. Long-range electrostatic interactions were calculated with the Particle-mesh Ewald (PME) summation scheme ^72^; Van der Waals and short-range Coulomb interactions were truncated at 10 Å.

### Structural Network Analysis with PyInteraph

We used the *PyInteraph* suite ^26^ to unveil the paths of long-range structural communication in Bcl-xL applying the principles of graph theory, focusing on the possible structural modifications which may be transmitted from the p53 binding site to the BH3-only proteins binding pocket upon binding of p53 and vice versa. We considered as interacting pairs any two residues whose side-chain centers of mass lied within 5.125 Å. This cut-off was selected on the basis of the results produced by the *PyInKnife* pipeline ^51^, which calculates the average number and size of the connected components and hubs distribution of a PSN implementing a Jackknife resampling method on the ensemble, estimating the variation of the data across the different resamplings. We tested contact cutoffs in the range of 5.0-5.5 Å, as suggested from other studies on globular proteins and different MD force fields ^50,51^. Specifically, we probed values every 0.125 Å. We noticed that a contact cut-off of 5.0 Å led to a quite sparse PSN with many poorly populated connected components (Fig. S1), whilst a tendency of the nodes to group in one unique connected component is observable already at 5.25 Å (Fig. S1). Since the two cutoffs described above underpin the construction of unrepresentative networks, we retained a contact cutoff of 5.125 Å for the analyses.

We also applied a cutoff to the persistence (p_crit_) of the interaction in our PSN to filter out transient and spurious interactions from our network. This cut-off was selected on the basis of the size of the largest connected component evaluated at different cut-offs (from 0-100% with step 10%), identifying a pcrit of 20% as previously observed for other cases of studies ^26^.

### Prediction of pathogenicity score with REVEL and collection of cancer-related mutations

REVEL is an ensemble method which accounts for 18 different pathogenicity prediction scores and trained with random forest model ^62^. Mutations with REVEL scores higher than 0.4 are predicted to be pathogenic, according to the original publication. Mutations with a REVEL score of 0 are those for which the predictor could not assign a score due to the fact that they do not account for single nucleotide changes and as a such not covered by this predictor. Missense mutations of Bcl-x_L_ found by genomics studies have been collected, aggregated from Cbioportal ^73^ and COSMIC ^74^.

### Structure-based estimation of the impact of mutations on protein stability

We employed the *FoldX* energy function from the *FoldX suite* ^75^ to carry out *in silico* saturation mutagenesis using a *Python* wrapper that we recently developed and previously applied to other case study ^50,76–78^. We collected an average ΔΔG for each mutation over the whole NMR ensemble of 20 conformers for both the free state of Bcl-x_L_ (PDB entry 2LPC) and its complex with PUMA (PDB entry 2M04). The ensemble was used to account for flexibility in the protein, as we recently did on another protein system ^50^ since *FoldX* energy function only allows for local conformational changes. We calculated the ΔΔG upon mutations associated to protein stability and binding of the interactor. We applied the *BuildModel* module from the FoldX suite and five independent runs for mutations in our scan. The typical prediction error of FoldX is about 0.8 kcal/mol ^79^. We then use twice the prediction error (i.e., 1.6 kcal/mol) as a threshold to discriminate between neutral and deleterious mutations in the analyses.

## Acknowledgments

This project was supported by the LEO foundation grant number LF17006 and the Carlsberg Distinguished Fellowship CF18-0314 to EP group and three Erasmus Plus for Traineeship Fellowships to VS to visit EP group in 2016, 2017 and 2018 by the University of Milano-Bicocca and the University of Bologna, respectively. The calculations described in this paper were performed with the support of the DeiC National Life Science Supercomputer at DTU. the EU PRACE-DECI GRANT 14th CancerBH3 and the ISCRA-Cineca HPC Grants HP10C8YXRK and HP10CBLBWO. The authors would like to thank Matteo Tiberti for technical help and useful suggestions.

---

Fig. S1 – Nodes distribution at different distance cut-offs Distribution of nodes in the five most populated connected components when the distance cut-off was set to a) 5 Å b) 5.125 Å c) 5.5 Å.

