## Supplementary figures and images for "Bcl-xL dynamics and cancer-associated mutations under the lens of protein structure network and biomolecular simulations"

### Supplementary Figure S1

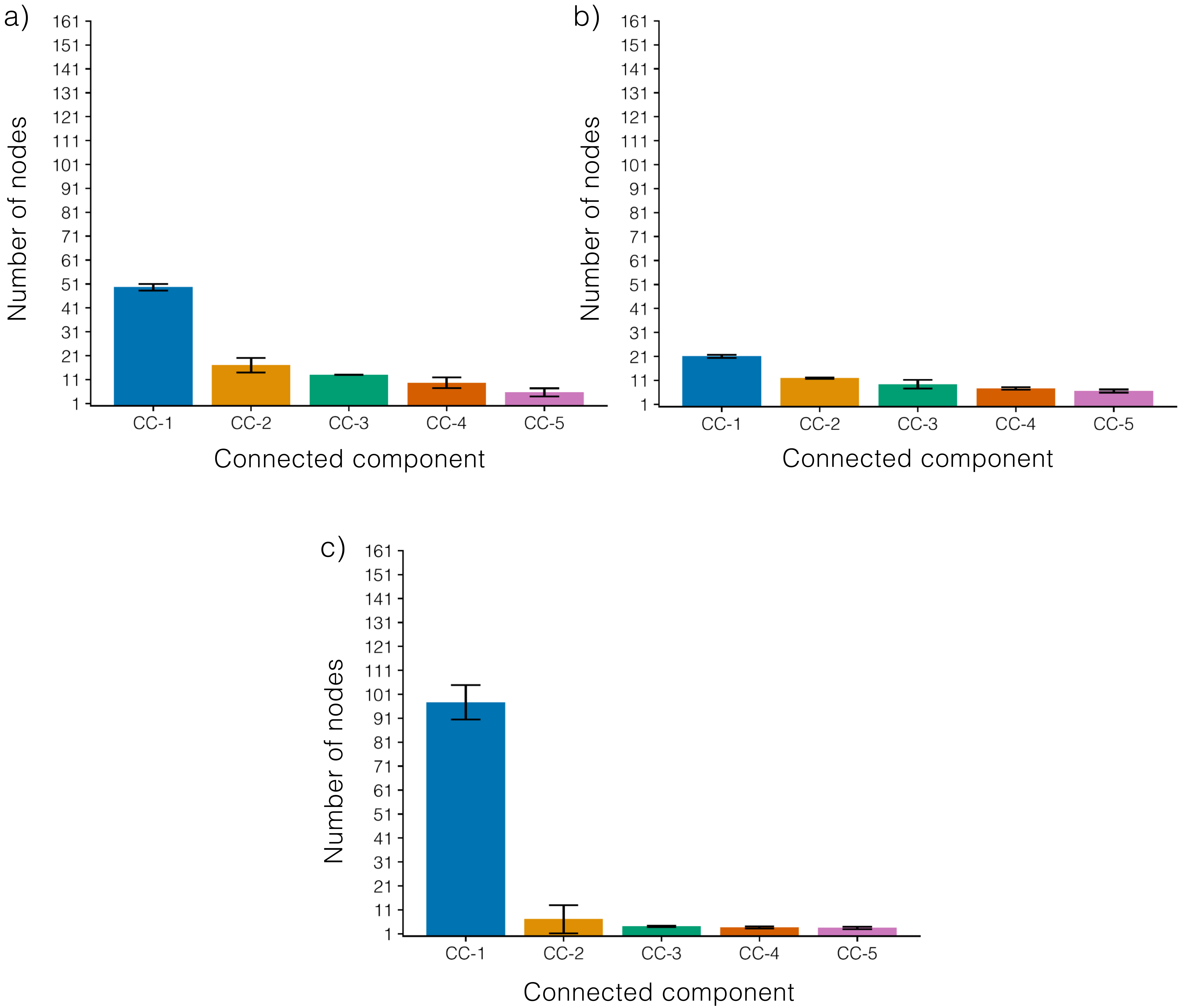

### Supplementary Text S1

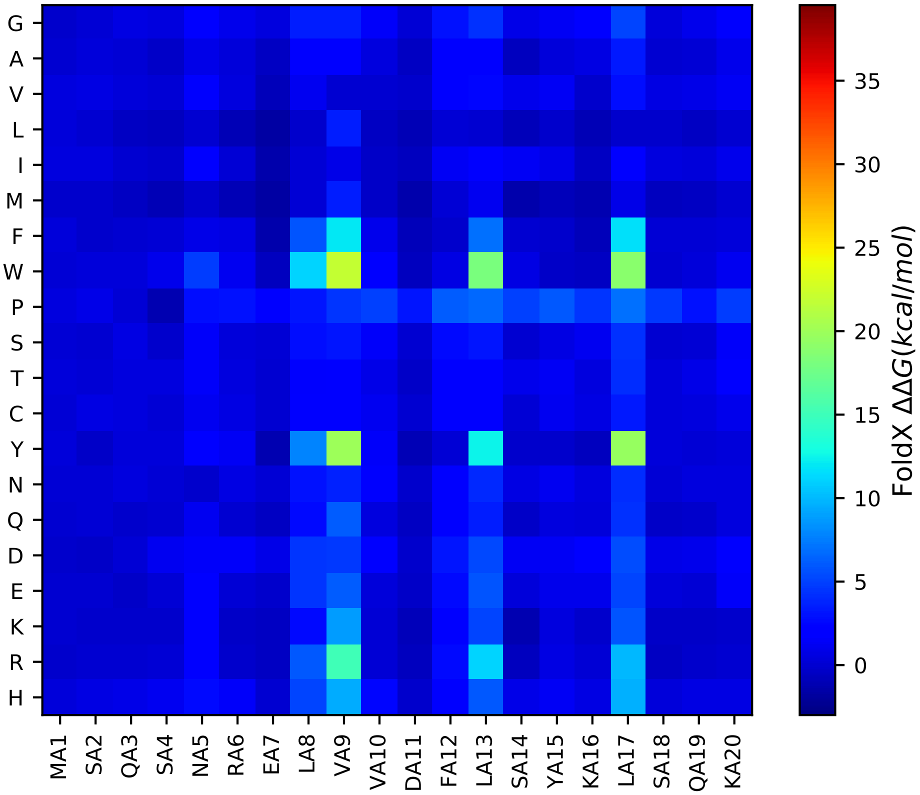

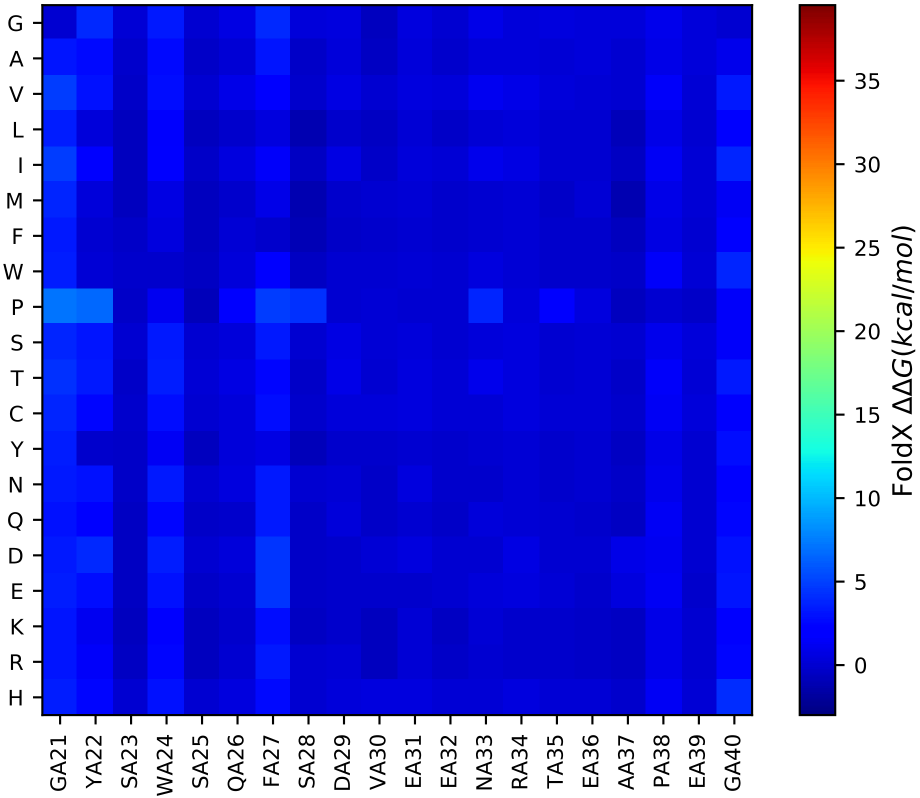

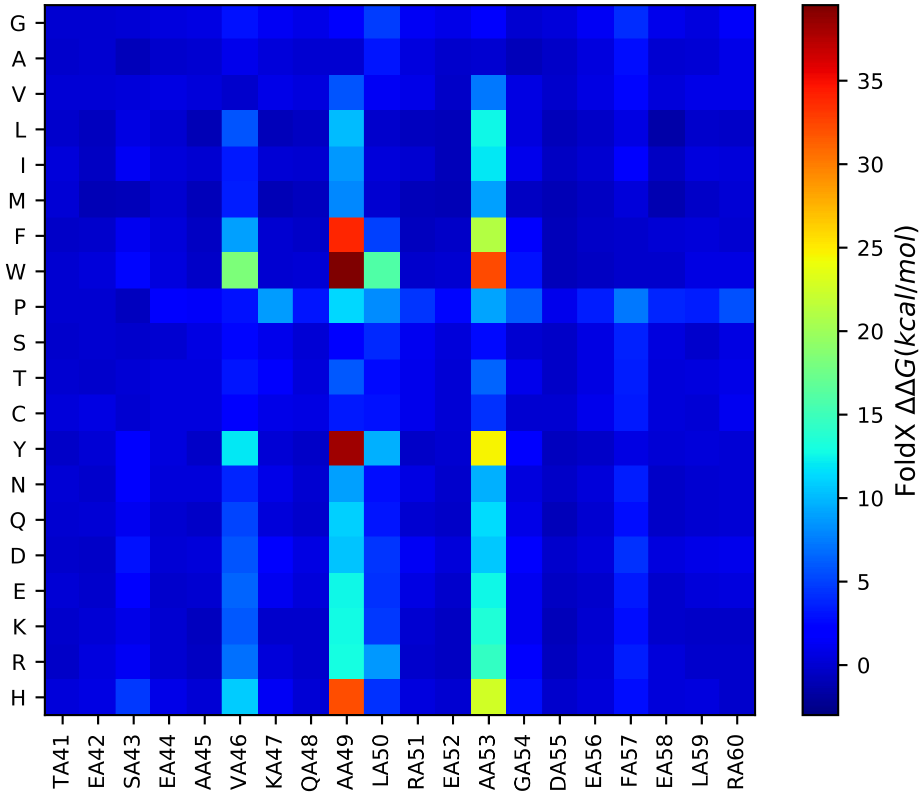

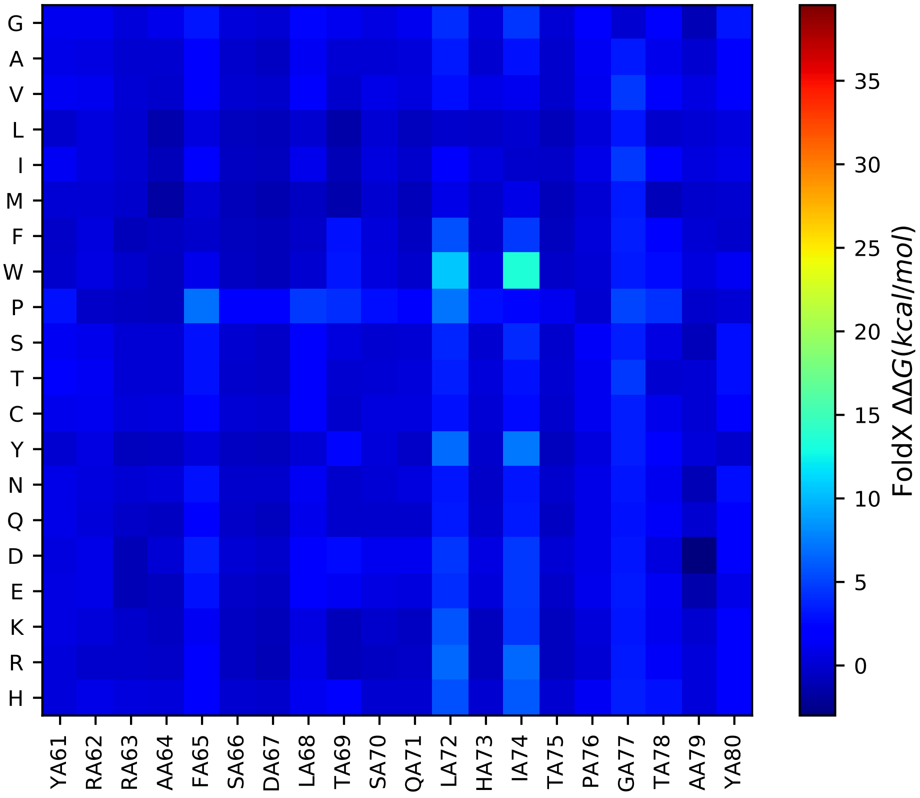

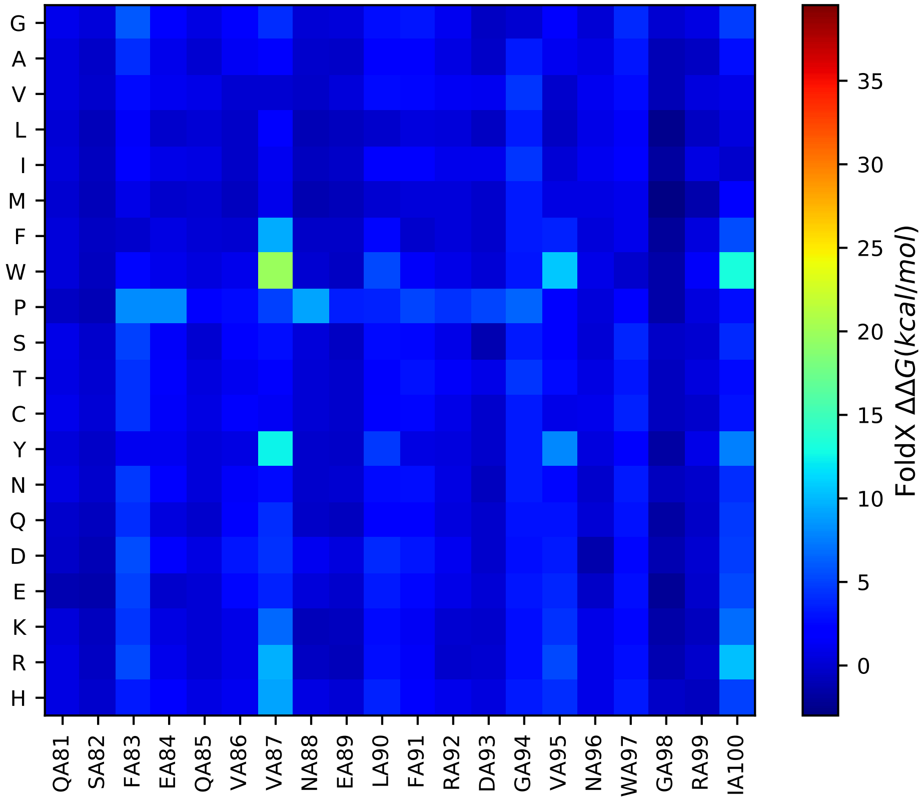

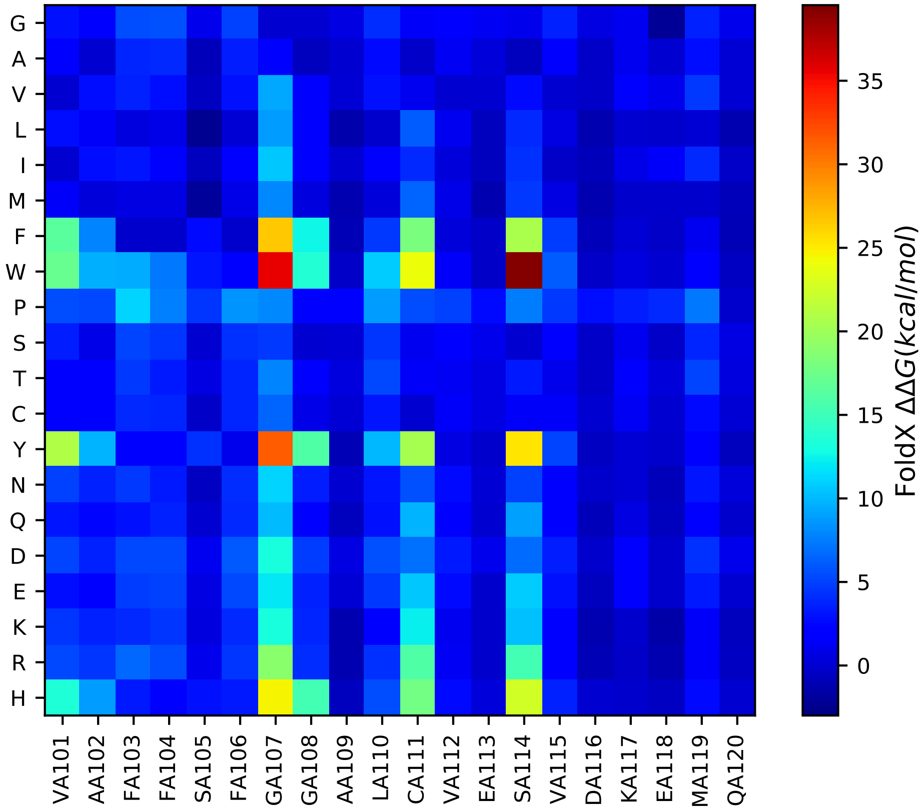

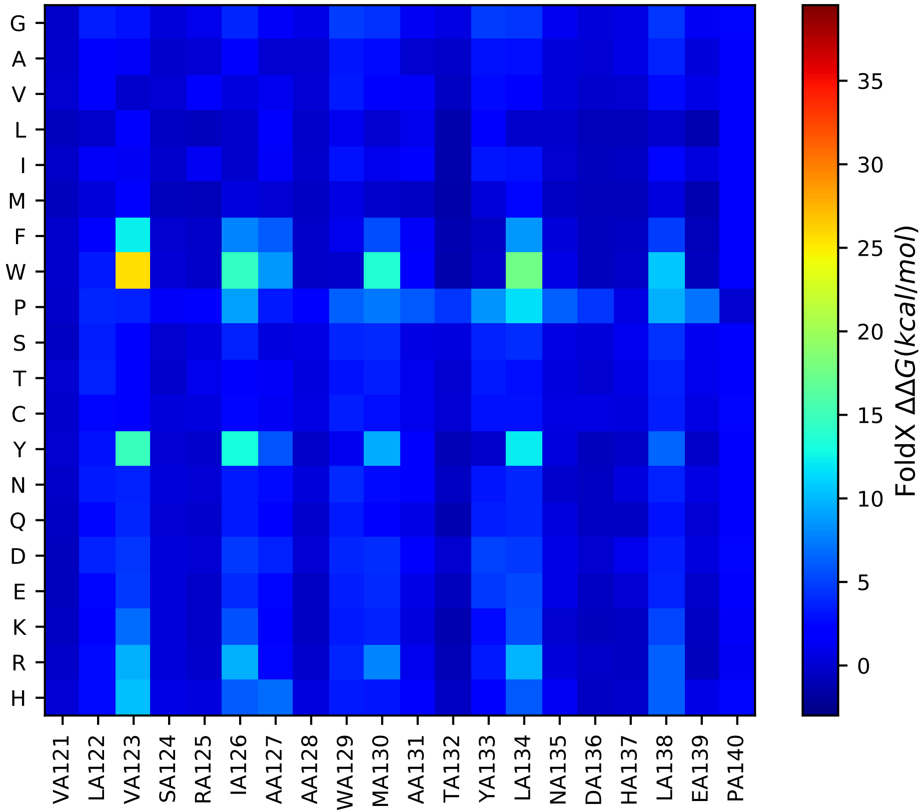

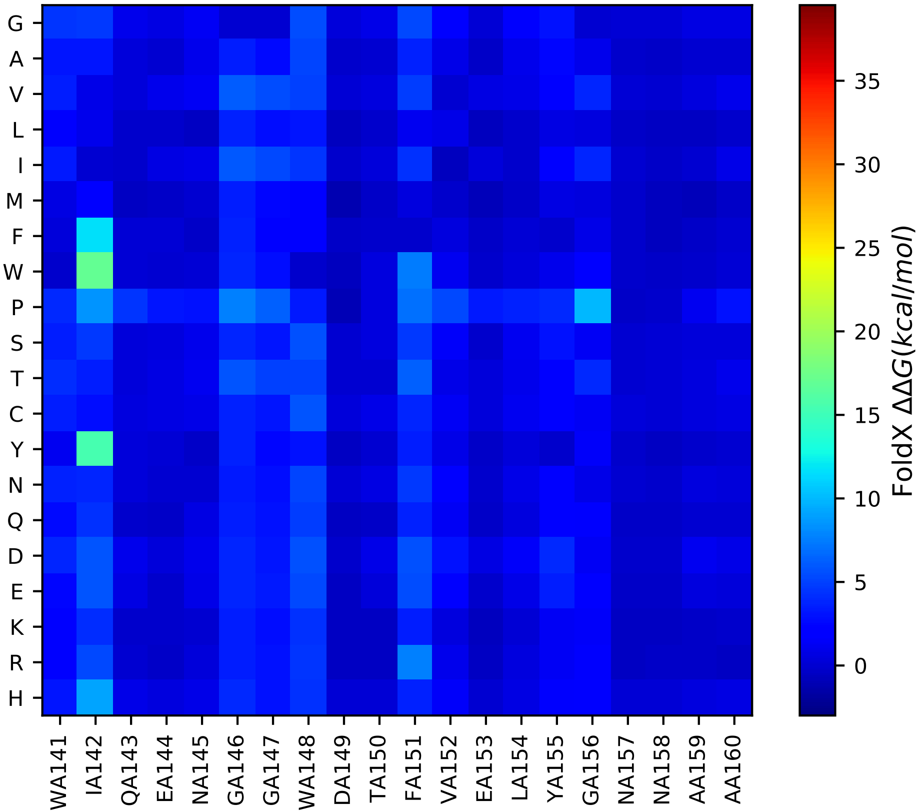

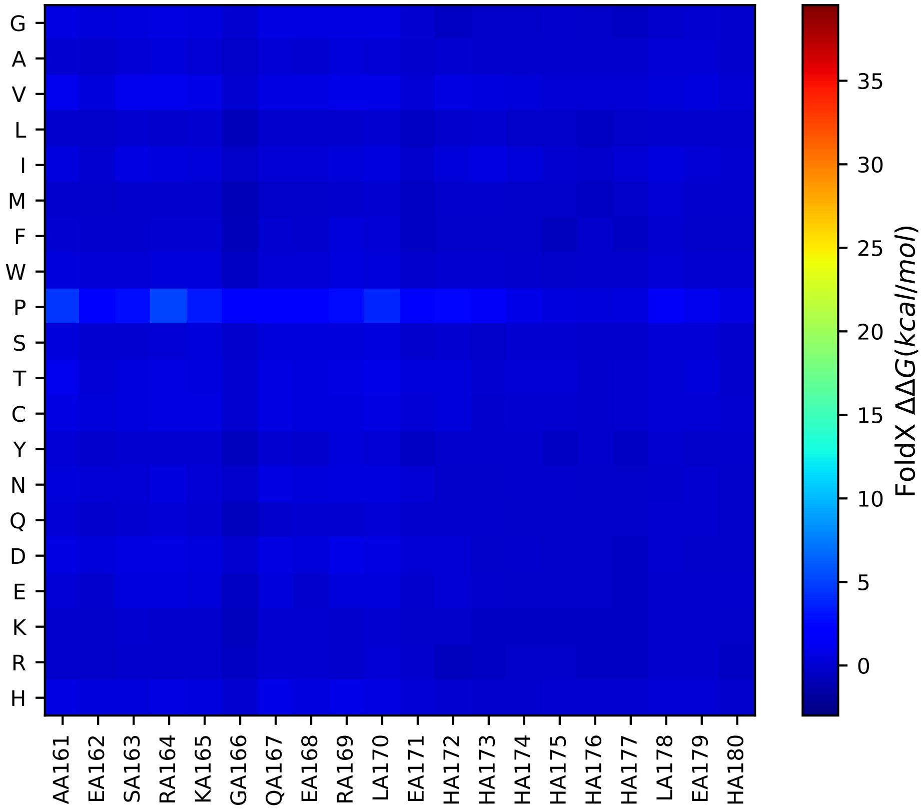

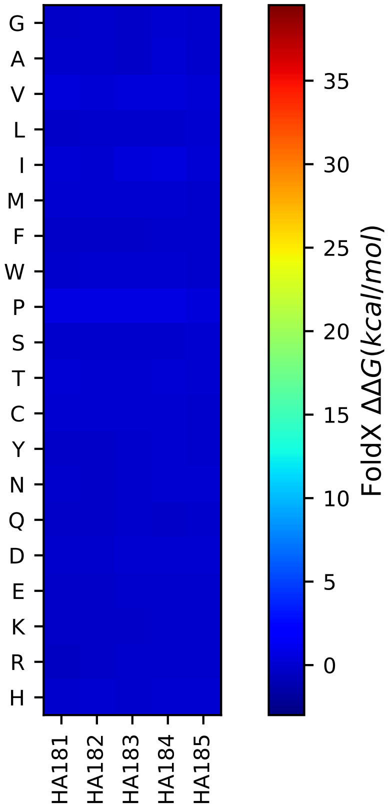

### Supplementary Text S2

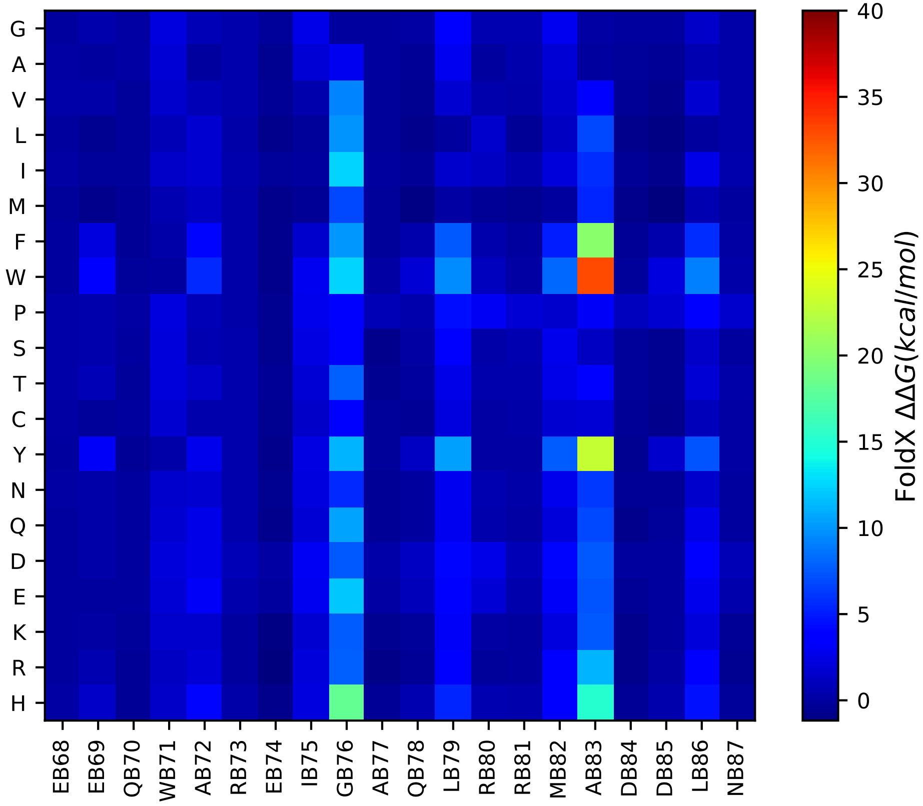

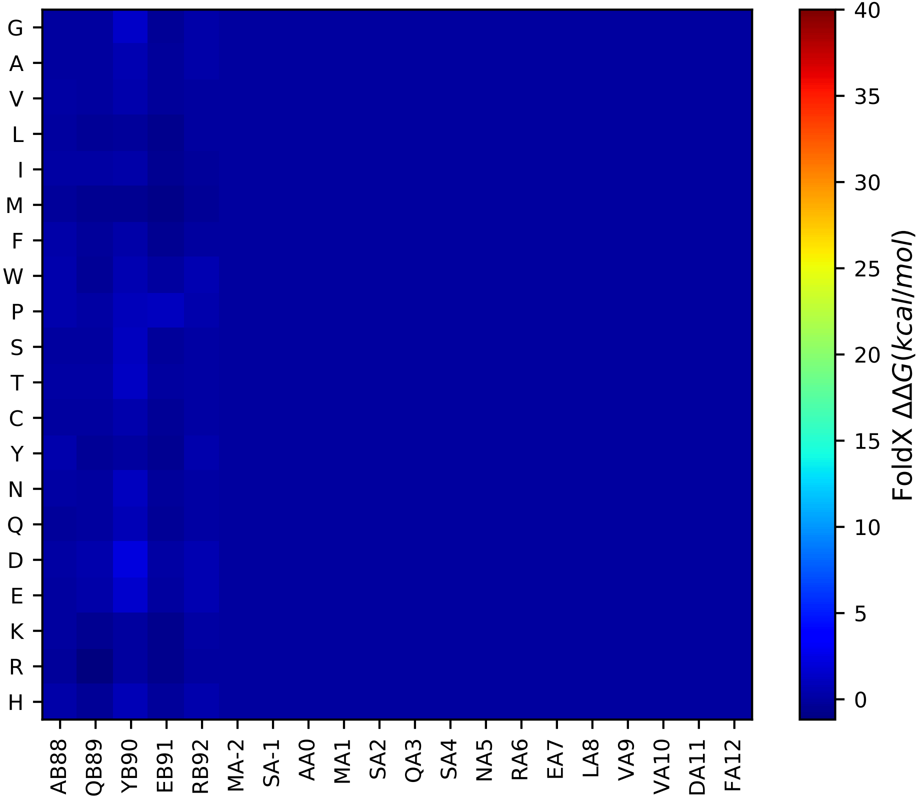

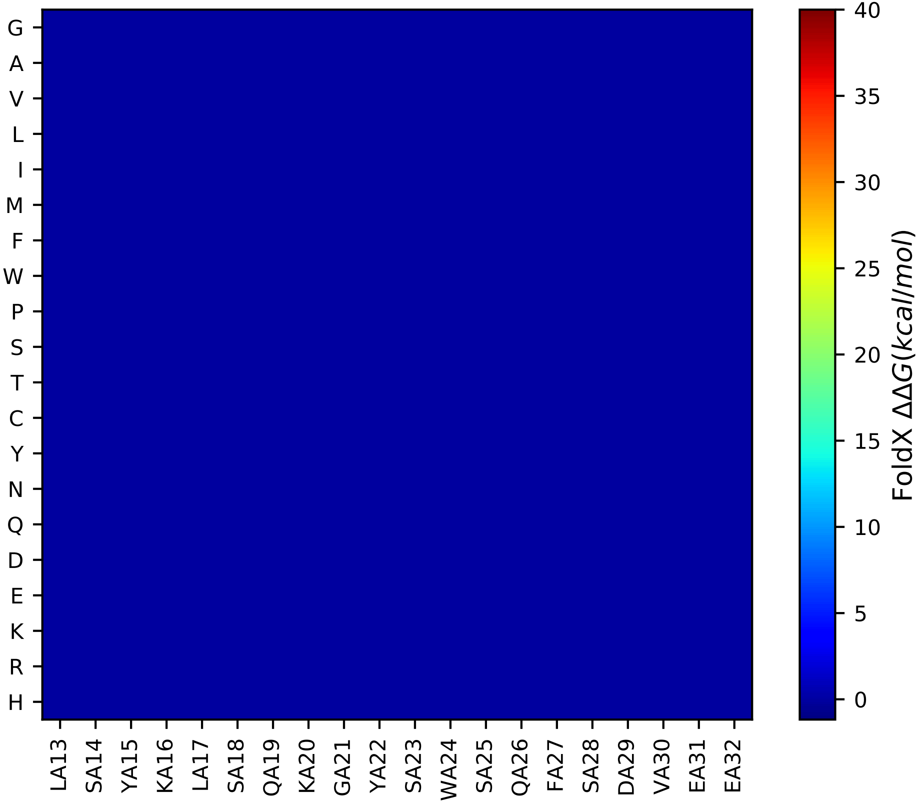

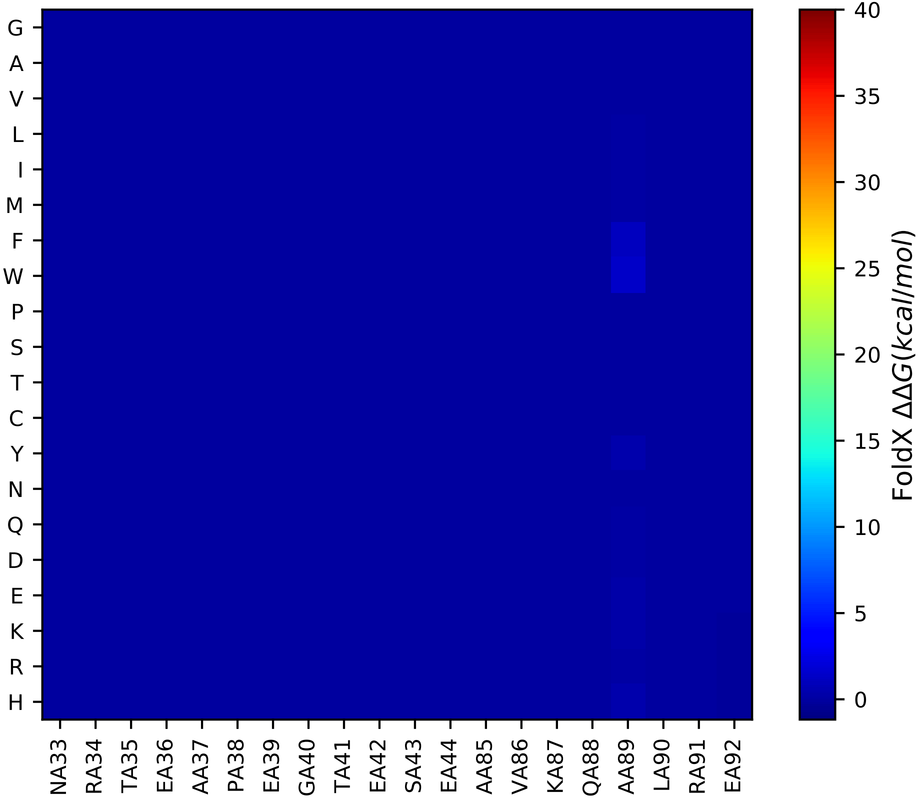

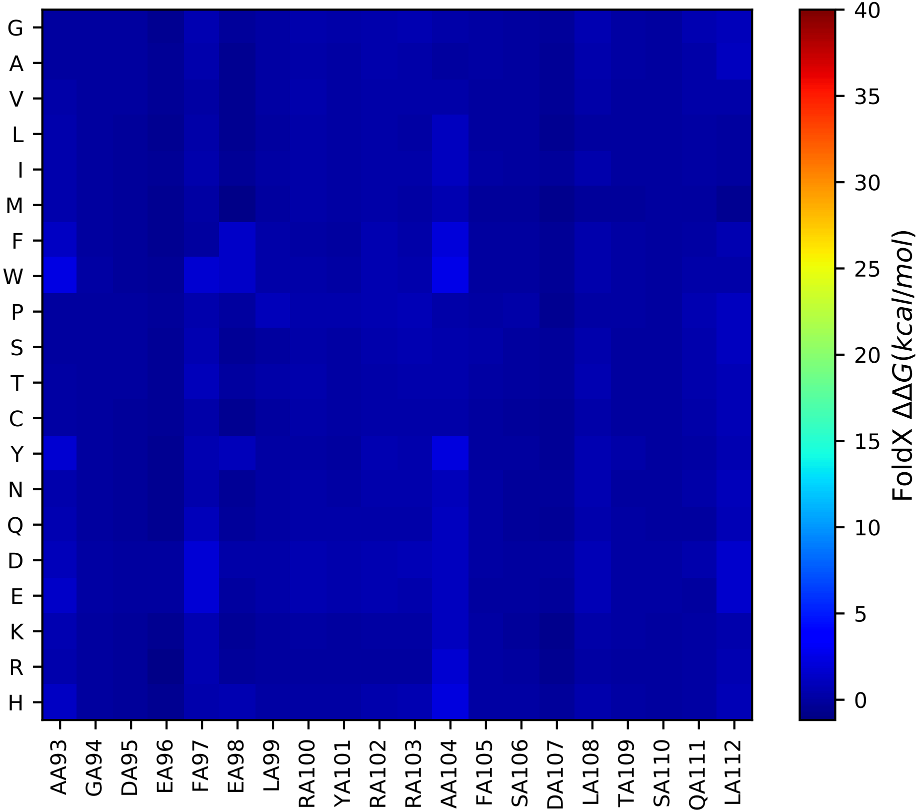

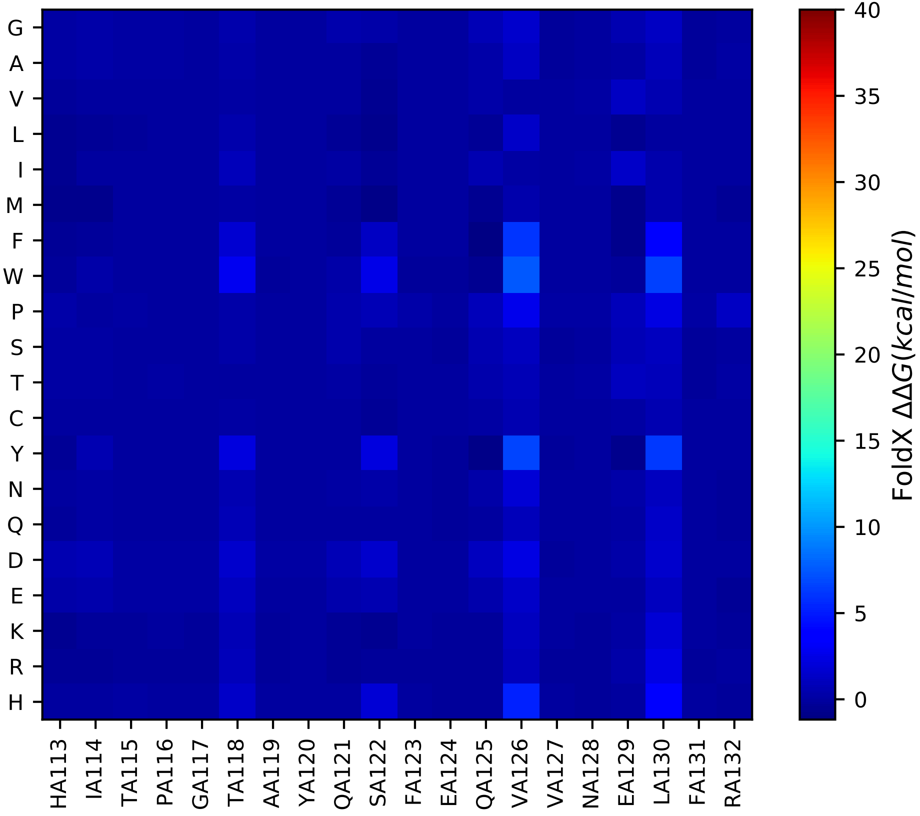

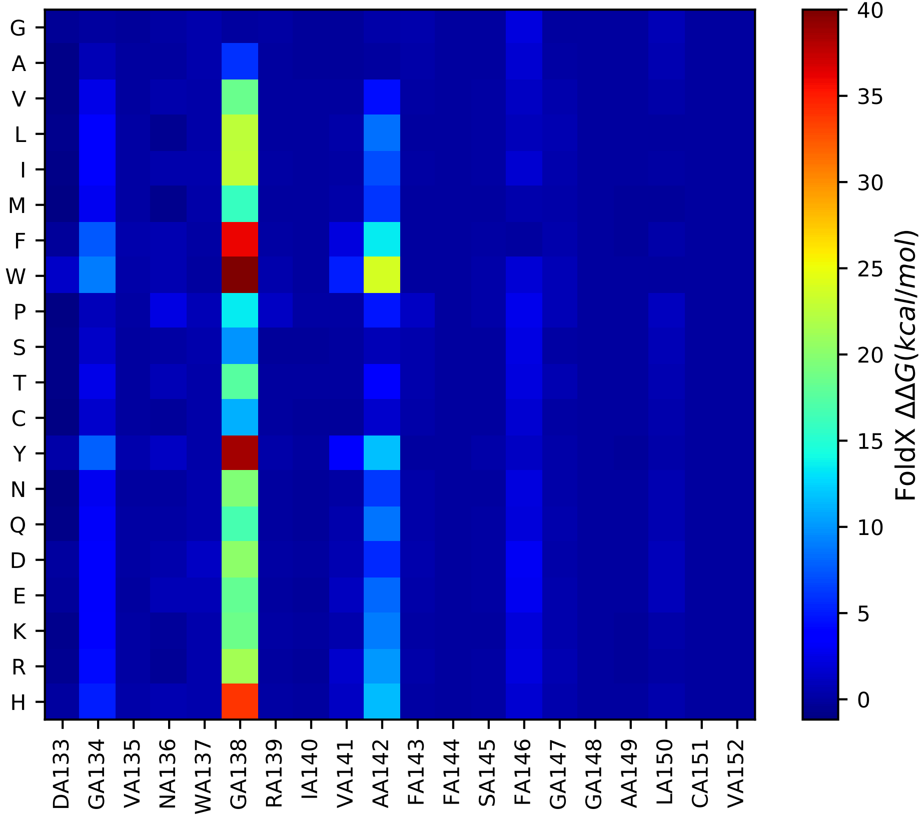

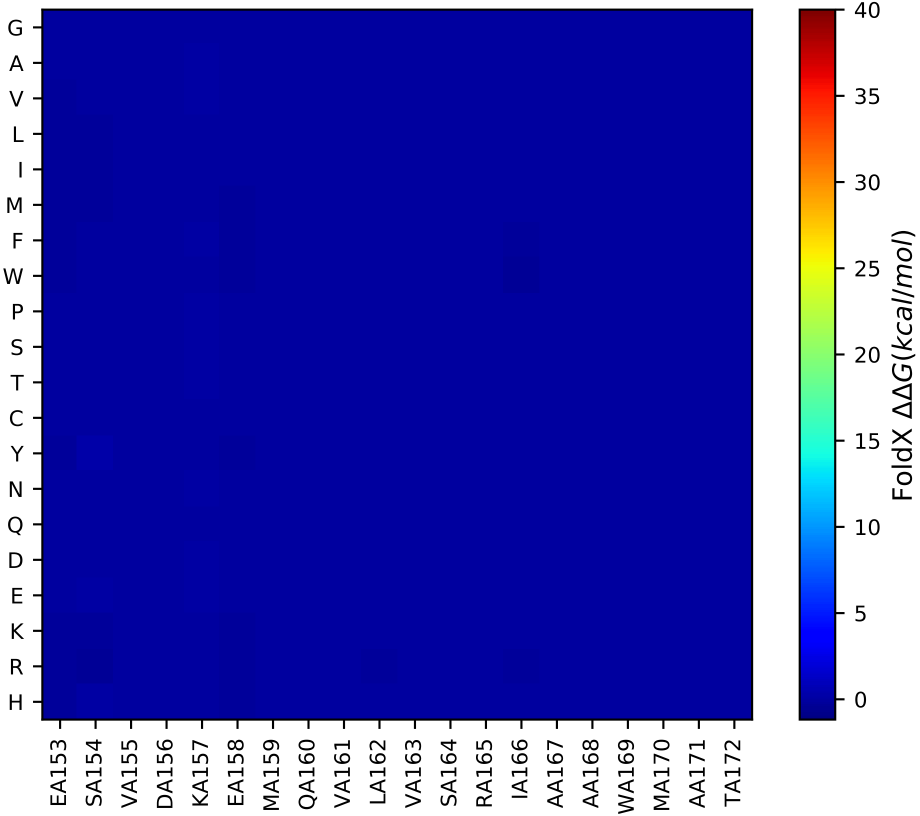

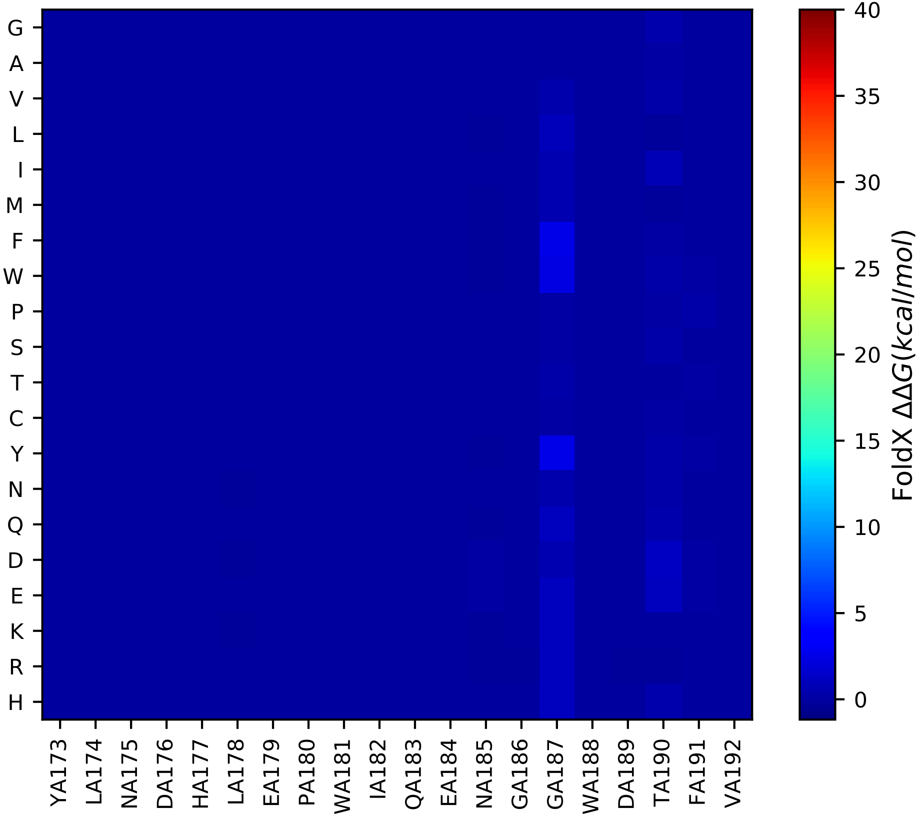

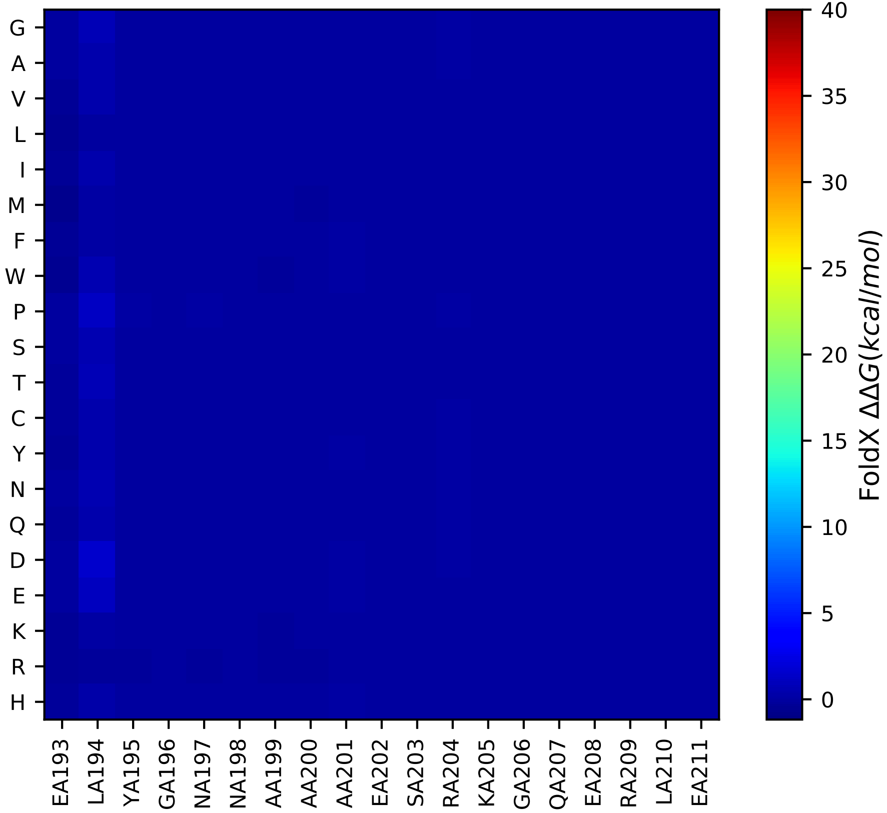
